# Unraveling the Core: Hub Genes Bridging Familial and Sporadic Parkinson’s Disease

**DOI:** 10.1101/2024.02.19.580936

**Authors:** Simran Singh, Gulshan Chauhan, Baisakhi Moharana, Sonia Verma

## Abstract

Parkinson’s disease (PD), a neurodegenerative disorder characterized by dopaminergic (DA) neuron loss in the substantia nigra, manifests as familial (genetically linked) or idiopathic (sporadic) forms. Despite distinct etiologies, both subtypes converge on shared pathological mechanisms that remain poorly understood. This study focused on identifying “hub genes” that might drive DA neuron degeneration in familial and idiopathic PD. The gene expression data from three publicly available datasets were reanalyzed. These datasets included samples from DA neurons derived from postmortem brains and patient-derived induced pluripotent stem cells. Twelve hub genes were identified to be dysregulated across all three datasets. The hub genes were found to play vital roles in membrane trafficking and vesicle-mediated transport. NetworkAnalyst-based reconstruction linked these hub genes to various other diseases. Experimental validation in neurotoxin-induced SH-SY5Y cell models of PD confirmed significant changes in the mRNA levels of some of the hub genes. Crucially, silencing one of the hub genes in *Caenorhabditis elegans* promoted DA neuron degeneration. Our study identifies potential candidates as therapeutic targets for DA neuron degeneration in familial and idiopathic PD.

## 1. Introduction

Parkinson’s disease (PD) is a multifaceted neurodegenerative disorder predominantly characterized by the progressive degeneration of dopaminergic (DA) neurons in the substantia nigra pars compacta, leading to hallmark motor deficits and non-motor symptoms ^1,2^. PD is broadly classified into two categories based on etiology: familial PD and idiopathic PD. Familial PD accounts for 5-10% of all cases and has been linked to mutations in pivotal genes such as *SNCA*, *LRRK2*, *PARK2*, *PINK1*, *DJ-1*, and *GBA*, each contributing to distinct cellular dysfunctions, including mitochondrial stress, impaired autophagy, proteostasis disruption, and lysosomal dysfunction ^3–18^. Genome-wide association studies (GWAS) have further identified novel risk loci such as *TMEM175*, *GAK*, and *BST1*, extending the genetic landscape of PD pathogenesis ^19,20^. Idiopathic PD accounts for approximately 90-95% of all PD cases ^21^. There are no clear genetic links in idiopathic cases, and individuals affected by this form of the disease typically lack a family history of PD. Understanding different forms of PD is crucial for developing targeted therapies for affected individuals.

Transcriptomic analyses have provided transformative insights into molecular mechanisms underlying PD. Studies have revealed dysregulated gene expression profiles in diverse sample types—including brain tissue, blood, serum exosomes, and skin—highlighting systemic molecular changes and “skin-brain” crosstalk. Notable findings include apoptosis-related gene expression alterations in the cingulate gyrus, copper metabolism-related hub genes in the substantia nigra (SN), heat shock protein dysregulation in blood samples, and complement system-associated hub genes in peripheral blood ^22–30^. Single-nucleus transcriptomics has illuminated selective cell vulnerability in the prefrontal cortex, while serum exosome profiling underscores intercellular communication as a potential biomarker and therapeutic target ^31,32^. Collectively, transcriptomic analyses across diverse sample types have paved the way for novel biomarkers, therapeutic targets, and a deeper understanding of the interplay between central and peripheral mechanisms in PD.

In both familial and idiopathic PD, the progressive loss of DA neurons located in SN is the defining feature of the disease. Despite the differences in their genetic basis, some common pathways and mechanisms may contribute to DA neuron degeneration in familial and idiopathic PD. Identifying the shared dysregulated genes and pathways can uncover core cellular processes contributing to DA neuron degeneration across PD forms. High-throughput RNA sequencing has been used to identify dysregulated genes by comparing gene expression patterns in healthy and PD-afflicted DA neurons ^33–39^. Samples such as postmortem brain tissue and induced pluripotent stem cells (iPSC)-derived models offer unique advantages; postmortem tissue provides invaluable insights into the actual pathological state of the disease, and iPSCs offer a dynamic and renewable platform for modeling disease progression and developing therapies ^33–39^. Together, these approaches enhance our understanding of PD and facilitate the development of novel therapies that could be effective for a broader population of PD patients, regardless of genetic background.

Several model organisms are employed to study PD ^40^. The invertebrate *Caenorhabditis elegans* presents unique advantages, including its small size, rapid lifecycle, and cost-effective maintenance, which enable high-throughput studies on large populations. Notably, *C. elegans* shares over 80% genetic homology with human disease-related genes and possesses DA neurons, making it an excellent platform for identifying and characterizing genes and factors involved in DA neuron degeneration ^41–44^. Moreover, human α-synuclein-expressing *C. elegans* models recapitulate key PD phenotypes, such as age-dependent α-synuclein aggregation, DA neuron loss, and disruptions in DA-dependent behaviors ^45–48^. Additionally, the neurotoxin 1-methyl-4-phenyl 1,2,3,6 tetrahydropyridine (MPTP)-treated SH-SY5Y is extensively employed to investigate the molecular and cellular mechanisms underlying PD pathology and to evaluate potential neuroprotective compounds for PD treatment ^49–55^. Together, *C. elegans* and SH-SY5Y cells offer robust and complementary tools for uncovering the mechanisms of PD and developing potential therapeutic interventions.

In this study, we integrated datasets from Novosadova, E. et al., Zaccaria, A. et al., and Kim, J.W. et al., representing RNA sequencing of DA neurons in familial and idiopathic PD (**Graphical Abstract**) ^34,35,39^. We identified 40 genes dysregulated in the DA neurons in all three databases. Analysis of the Protein-Protein Interaction (PPI) network of these genes identified 12 hub genes. Gene ontology (GO) analysis highlighted the functional roles of genes in various biological processes, molecular functions, and cellular components. Using the Reactome database, we identified enriched pathways in which the genes participate, shedding light on fundamental cellular processes related to DA neuron degeneration in PD. Our findings were validated in the *Caenorhabditis elegans* and MPTP-induced SH-SY5Y model of PD. These findings underscore the importance of shared molecular genes and pathways in PD pathogenesis and highlight dysregulated hub genes as promising candidates for therapeutic intervention.

## 2. Material and Methods

### 2.1 *In-silico* Experiments

#### 2.1.1. Data Extraction

The data used in this study were obtained from the Gene Expression Omnibus (GEO) (http://www.ncbi.nlm.nih.gov/geo/) database ^56^. The GEO database is a publicly accessible resource for functional genomic data, including microarrays, chips, and high-throughput gene expression datasets. The keyword “Parkinson’s Disease” was used to search the database, and the search results were filtered for the Organism “*Homo sapiens*” and Study type “Expression profiling by high throughput sequencing.” Among the filtered results, datasets meeting the following criteria were included:

1. Profiling conducted on:
  a. SN-DA neurons from post-mortem brains, or
  b. Human iPSC-derived DA neurons
2. Samples included PD and healthy controls.
3. Analysis with GEO2R is available.

Based on the inclusion criteria, three gene datasets (GSE90469, GSE169755, and GSE181029) were selected for further analysis ^34,35,39^. Further information on these datasets is summarized in **Table 1**.

**Table 1:**
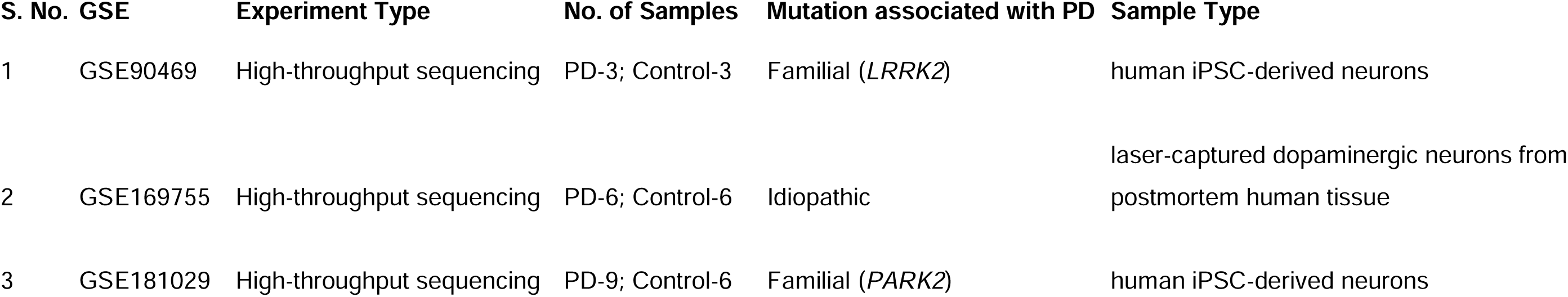
GEO studies included in the analysis.

| **S. No.** | **GSE** | **Experiment Type** | **No. of Samples** | **Mutation associated with PD** | **Sample Type** |
| --- | --- | --- | --- | --- | --- |
| 1 | GSE90469 | High-throughput sequencing | PD-3; Control-3 | Familial (*LRRK2*) | human iPSC-derived neurons |
| 2 | GSE169755 | High-throughput sequencing | PD-6; Control-6 | Idiopathic | laser-captured dopaminergic neurons from postmortem human tissue |
| 3 | GSE181029 | High-throughput sequencing | PD-9; Control-6 | Familial (*PARK2*) | human iPSC-derived neurons |

#### 2.1.2. Differentially Expressed Genes (DEGs) Identification

GEO2R, an online sequencing data analysis tool developed by the GEO database, was used to analyze gene expression profiles and identify DEGs between PD and control DA neurons for each dataset. GEO2R utilizes two R packages (GEOquery and Limma) for its analyses (https://www.r-project.org) ^57–59^. GEOquery reads the data, while Limma calculates differential expressions. The samples used to perform analysis in each dataset are summarised in **Table 2**. The *p* ≤ 0.05 and |log_2_ Fold-Change (FC)| ≥ 0.5 were set as thresholds for DEGs. Gene expression distributions for each dataset were visualized using volcano plots and box plots. Overlapping DEGs from the three datasets were defined as common DEGs (cDEGs) and displayed using Venn diagrams created with the online platform Venny 2.1.0 (http://bioinfogp.cnb.csic.es/tools/venny/index.html).

**Table 2:**
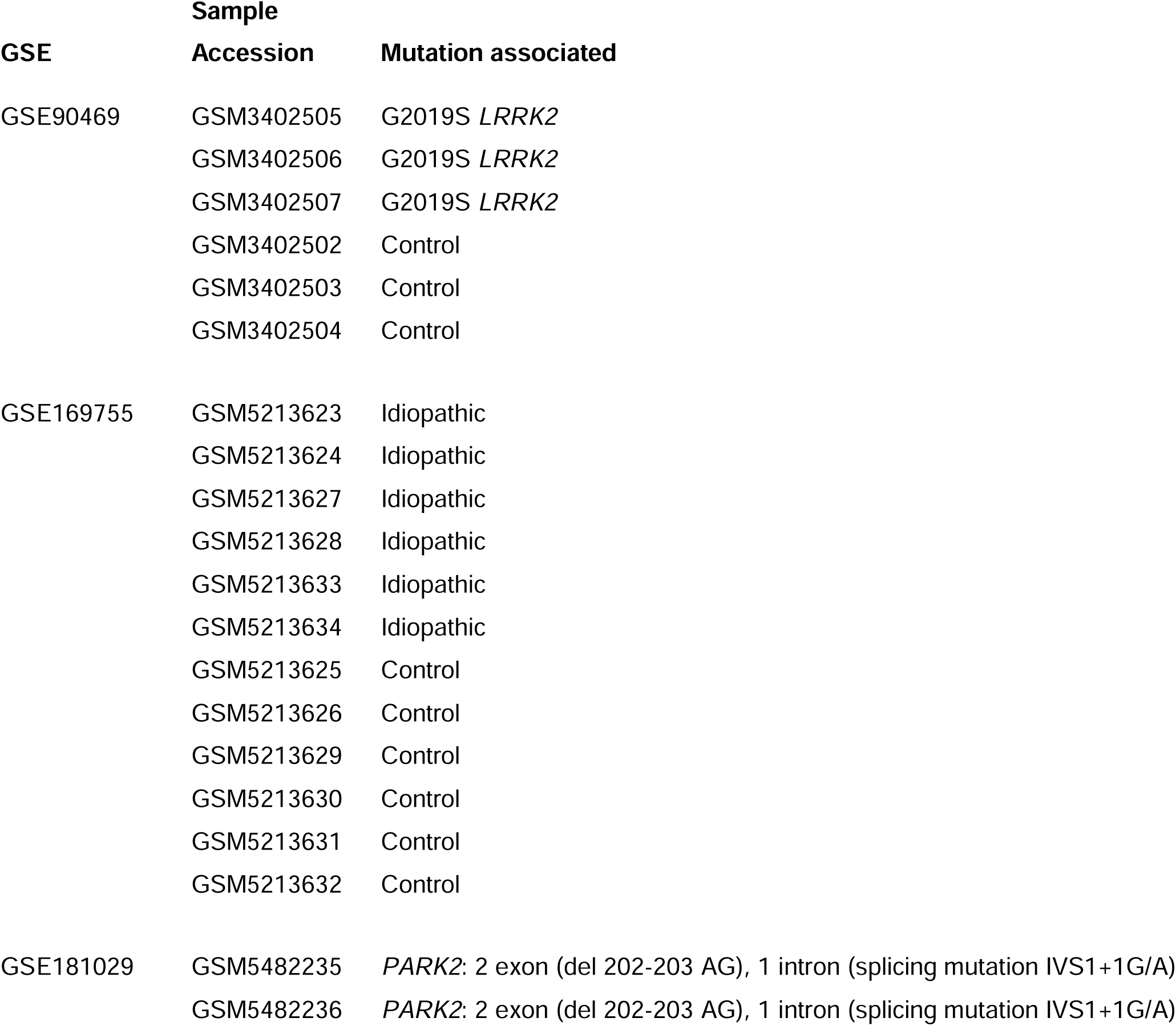
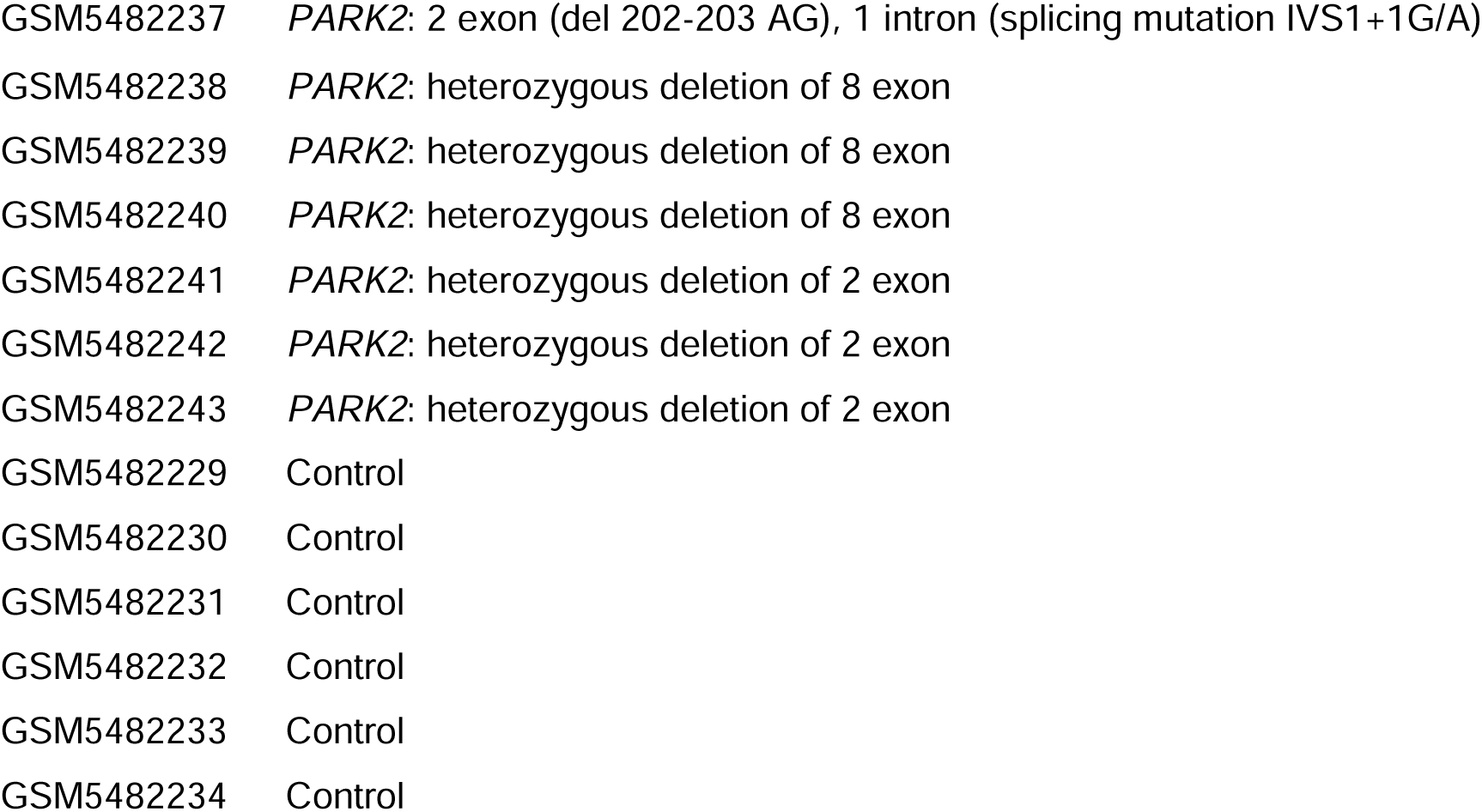
Overview of samples used for GEO2R analysis in each dataset.

| GSE | Sample |  |
| --- | --- | --- |
|  | Accession | Mutation associated |
| GSE90469 | GSM3402505 | G2019S *LRRK2* |
|  | GSM3402506 | G2019S *LRRK2* |
|  | GSM3402507 | G2019S *LRRK2* |
|  | GSM3402502 | Control |
|  | GSM3402503 | Control |
|  | GSM3402504 | Control |
| GSE169755 | GSM5213623 | Idiopathic |
|  | GSM5213624 | Idiopathic |
|  | GSM5213627 | Idiopathic |
|  | GSM5213628 | Idiopathic |
|  | GSM5213633 | Idiopathic |
|  | GSM5213634 | Idiopathic |
|  | GSM5213625 | Control |
|  | GSM5213626 | Control |
|  | GSM5213629 | Control |
|  | GSM5213630 | Control |
|  | GSM5213631 | Control |
|  | GSM5213632 | Control |
| GSE181029 | GSM5482235 | *PARK2*: 2 exon (del 202-203 AG), 1 intron (splicing mutation IVS1+1G/A) |
|  | GSM5482236 | *PARK2*: 2 exon (del 202-203 AG), 1 intron (splicing mutation IVS1+1G/A) |

|  |  |
| --- | --- |
| GSM5482237 | *PARK2*: 2 exon (del 202-203 AG), 1 intron (splicing mutation IVS1+1G/A) |
| GSM5482238 | *PARK2*: heterozygous deletion of 8 exon |
| GSM5482239 | *PARK2*: heterozygous deletion of 8 exon |
| GSM5482240 | *PARK2*: heterozygous deletion of 8 exon |
| GSM5482241 | *PARK2*: heterozygous deletion of 2 exon |
| GSM5482242 | *PARK2*: heterozygous deletion of 2 exon |
| GSM5482243 | *PARK2*: heterozygous deletion of 2 exon |
| GSM5482229 | Control |
| GSM5482230 | Control |
| GSM5482231 | Control |
| GSM5482232 | Control |
| GSM5482233 | Control |
| GSM5482234 | Control |

#### 2.1.3. PPI Network Analysis

A PPI network of the proteins encoded by the cDEGs was predicted and constructed using the GeneMANIA plugin of Cytoscape software (version: 3.8.0; http://cyto-scape.org/) ^60^. The minimum required interaction score for the PPI analysis was set at a combined score ≥ 0.4. Nodes represented proteins, while edges denoted interactions between proteins. The Molecular Complex Detection (MCODE) plugin in Cytoscape was employed to perform network clustering analysis to identify key functional modules. These modules represent groups of highly interconnected proteins within the PPI network, suggesting they may participate in related biological pathways critical to DA neuron function and survival. The selection criteria for clustering included K-core = 2, degree cutoff = 2, max depth = 100, and node score cutoff = 0.2. Significant modules identified through MCODE analysis were used to identify hub genes.

#### 2.1.4. Functional Enrichment Analysis

Functional enrichment analysis of GO terms was conducted using the Database for Annotation, Visualization, and Integrated Discovery (DAVID) (https://david.ncifcrf.gov/) ^61^. Significantly enriched function annotations were defined as GO terms and Reactome pathways with *p* ≤ 0.05. GO analysis included biological processes, cellular components, and molecular functions to describe the activity and localization of genes. The bubble plots of GO terms and Reactome pathways were visualized via the “ggplot2” package of RStudio software (version 1.3.959; https://rstudio.com/) ^62^.

#### 2.1.5. Gene-Disease Association Analysis

Diseases often share genetic underpinnings, with some genes implicated in multiple disorders. Understanding these gene-disease associations can provide insights into the broader implications of gene dysregulation beyond PD. Gene-disease associations were studied using the DisGeNET database integrated into the NetworkAnalyst platform. DisGeNET aggregates data from sources like GWAS catalogs and literature collections, facilitating insights into genetic underpinnings and comorbidities of diseases ^63^.

### 2.2. Cell-culture Experiments

#### 2.2.1. Cell Culture Maintenance

The human neuroblastoma cell line SH-SY5Y (ATCC) was cultured in Dulbecco’s Modified Eagle Medium (Gibco™) supplemented with 10% fetal bovine serum (Gibco™) and 1% penicillin-streptomycin (Gibco™). Cells were maintained in a humidified incubator at 37°C and 5% CO_2_ ^64^.

#### 2.2.2. MPTP Treatment

A 10 mM stock solution of MPTP (Sigma-Aldrich) was prepared in dimethyl sulfoxide (DMSO; Sigma-Aldrich). When the cells were 80% confluent, 10^6^ cells/ well were added to a 6-well plate. MPTP was diluted in cell media, and the cells were exposed to MPTP at a final concentration of 200 μM for 24 h. Cells treated with 0.1% DMSO were used as control ^65^.

#### 2.2.3. RNA Isolation and Quantitative Reverse Transcription Polymerase Chain Reaction (qRT-PCR)

RNA was isolated using the RNeasy Mini Kit (Qiagen). About 2μg of RNA was reverse-transcribed into cDNA with the cDNA Reverse Transcription Kit (Promega). qRT-PCR was performed using SYBR Premix Ex Taq^TM^ (TaKaRa Bio Inc.) and RT-PCR machine (Biorad-CFX96). Gene expression levels were normalized to *GAPDH* and evaluated using the 2^−ΔΔCt^ method. All the primers used for the analysis are listed in **Supplementary Table 1**.

### 2.3. *C. elegans* Experiments

#### 2.3.1. *C. elegans* Strains and Maintenance

The strains BY250 (P*_dat-1_*::GFP) and UA44 (P*_dat-1_*::GFP, P*_dat-1_*:: human-α-synuclein^wt^) were procured from *Caenorhabditis Genetics Center* (CGC), University of Minnesota, MN, USA, and maintained on the nematode growth medium (NGM) plate seeded with *Escherichia. coli* OP50 as the maintenance food at 20°C ^66^. Adult worms were treated with a bleaching solution to isolate eggs, which were then age-synchronized at the first larval stage (L1) in M9 buffer at 20°C for 16-18 h ^66^. The synchronized L1 larvae were subsequently transferred to NGM RNA Interference (RNAi) plates for gene knockdown experiments.

#### 2.3.2. RNAi Induction

RNAi was performed by feeding worms with *E. coli* strain HT115 (DE3) carrying a plasmid (L4440) expressing double-stranded RNA targeting a gene of interest ^67^. HT115 (DE3) carrying the empty vector was used as a control RNAi. The bacteria were cultured overnight in Luria-Bertani broth (LB) containing 100 μg/ml ampicillin (Sigma) and 12.5 µg/ml tetracycline (Sigma) at 37°C. This overnight culture was used to inoculate fresh LB containing 100 µg/ml ampicillin in a 1:100 ratio. The secondary cultures were grown at 37°C until they reached an OD_600_ of 0.4. The cultures were then centrifuged, and the pellets were resuspended in M9 buffer containing 1mM isopropyl β-D-1-thiogalactopyranoside (IPTG) (Sigma) and 100 µg/ml ampicillin. 400 μl of the resuspended culture was used to plate 60 mm NGM plates supplemented with 100 μg/ml ampicillin and 2mM IPTG. The seeded plates were left in a laminar hood for 16 h and then stored at 4°C until L1-stage worms were transferred onto the RNAi plates. To prevent progeny growth, 40 μM fluoro-2’-deoxy-β-uridine (TCI) was added at the fourth larval stage ^68^.

#### 2.3.3. DA Neuron Imaging

Three-day-old adult worms were collected from RNAi plates, washed with M9 buffer, and transferred onto a 2% agarose pad on a glass slide. The worms were anesthetized with 30 mM sodium azide (Sigma) and covered with a coverslip. Images of the head region were captured at 20X magnification using a fluorescence microscope (Carl Zeiss), and the fluorescence intensity of GFP in DA neurons was quantified using ImageJ software (NIH, Bethesda, MD) ^69^. A minimum of 20 worms were imaged for each RNAi experiment.

#### 2.3.4. RNA Isolation and qRT-PCR

Worms at day three of adulthood were collected and washed with M9 buffer. Trizol reagent (Invitrogen™) was added at approximately four times the volume of the worm pellet, and the worms were lysed through two freeze-thaw cycles followed by vigorous vortexing. RNA was purified using phenol:chloroform: isoamyl alcohol extraction and isopropanol precipitation ^70^. qRT-PCR was performed following the protocol described in section 2.2.4, with gene expression levels normalized to β-actin and analyzed using the 2^−ΔΔCt^ method ^70^. The primers used for the analysis are listed in **Supplementary Table 1**.

### 2.4. Statistical Analysis of Cells and *C. elegans* Experiments

Statistical analysis was conducted using GraphPad Prism 8 software (GraphPad, Inc., La Jolla, CA, United States). Data were expressed as mean ± standard deviation, and statistical significance was evaluated using Student’s t-test, with p ≤ 0.05 considered statistically significant.

## 3. Results

### 3.1. Identification of cDEGs among the datasets GSE90469, GSE169755, and GSE181029

GEO2R analysis of GSE90469 identified 1,384 DEGs (p ≤ 0.05 and |log_2_FC| > 0.5), of which 611 were upregulated, and 773 were downregulated in iPSC-derived DA neurons from patients with *LRRK2* (G2019S)-PD compared to controls (**Figure 1A**). In GSE169755, the analysis revealed 1,995 DEGs (p ≤ 0.05 and |log_2_FC| > 0.5), of which 1,234 were upregulated, and 761 were downregulated in DA neurons isolated from the SN of idiopathic PD patients as compared to controls (**Figure 1A**). From the GSE181029 dataset, 5,501 DEGs (p ≤ 0.05 and |log_2_FC| > 0.5) were identified, of which 2,843 were upregulated, and 2,658 were downregulated in iPSC-derived DA neurons from patients with three different *PARK2* mutations (**Figure 1A**). Gradient volcano plots illustrating the distribution of DEGs in each dataset are shown in **Figure 1B**. A Venn diagram revealed that 21 genes were upregulated and 19 were down-regulated in all three datasets (Fig. 1C-E). The identification of 40 cDEGs suggests a mechanistic contribution to PD pathogenesis, highlighting candidates for in-depth mechanistic studies into neurodegenerative processes.

**Figure 1.**
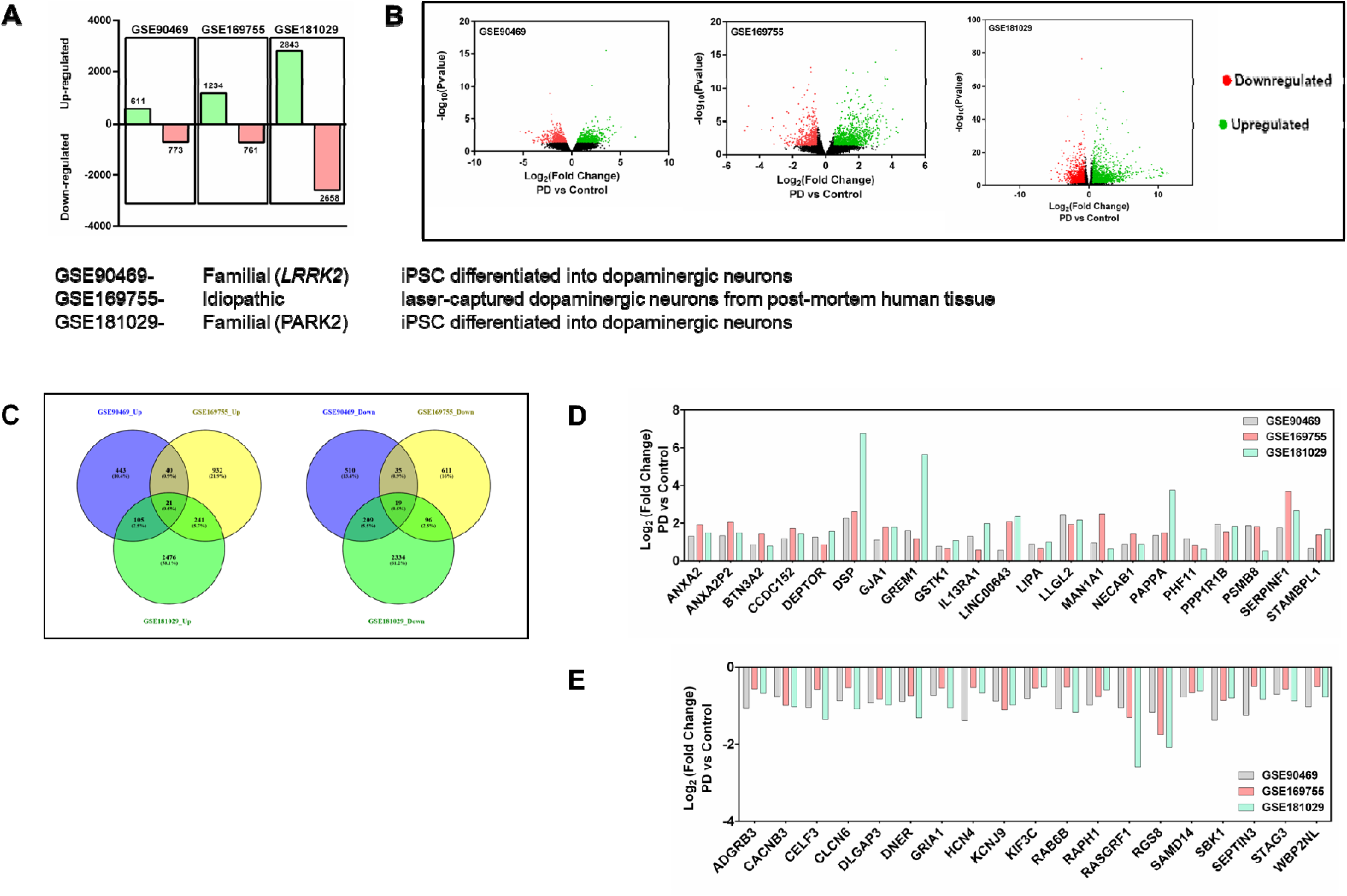
Gene expression analysis of the three selected datasets. (A) Bar graph of up- and downregulated genes in PD vs healthy DA neurons from each dataset. (B) Volcano plots to demonstrate gene expression differences between PD and control groups in the datasets. The criteria for selection of differentially expressed genes was set p ≤ 0.05 and |log_2_FC| ≥ 0.5. (C) In all three datasets, 21 genes were upregulated, and 19 genes were downregulated, as depicted in the Venn diagram. (D) Bar Graph showing the log_2_FC of upregulated genes in PD vs. Control DA neurons in respective datasets. (E) Bar Graph showing the log_2_FC of downregulated genes in PD vs. Control DA neurons in respective datasets.

### 3.2. Generation of PPI Network of cDEGs

The PPI network of cDEGs, constructed using Cytoscape based on the GeneMANIA database, included 58 nodes and 326 edges (**Figure 2A**). The MCODE plugin was applied to identify significant clusters or “modules” within the network. The analysis revealed three significant modules: Module 1 contained 17 nodes and 71 edges, Module 2 contained four nodes and nine edges, and Module 3 contained three nodes and six edges (**Figure 2B**). The nodes in these three modules included 12 cDEGs-*CELF3, PHF11, RAB6B, ADGRB3, BTN3A2, GRIA1, HCN4, KIF3C, PSMB8, RGS8, SEPTIN3,* and *SAMD14*. These 12 cDEGs were identified as hub genes, representing the most highly interconnected nodes within the network.

**Figure 2.**
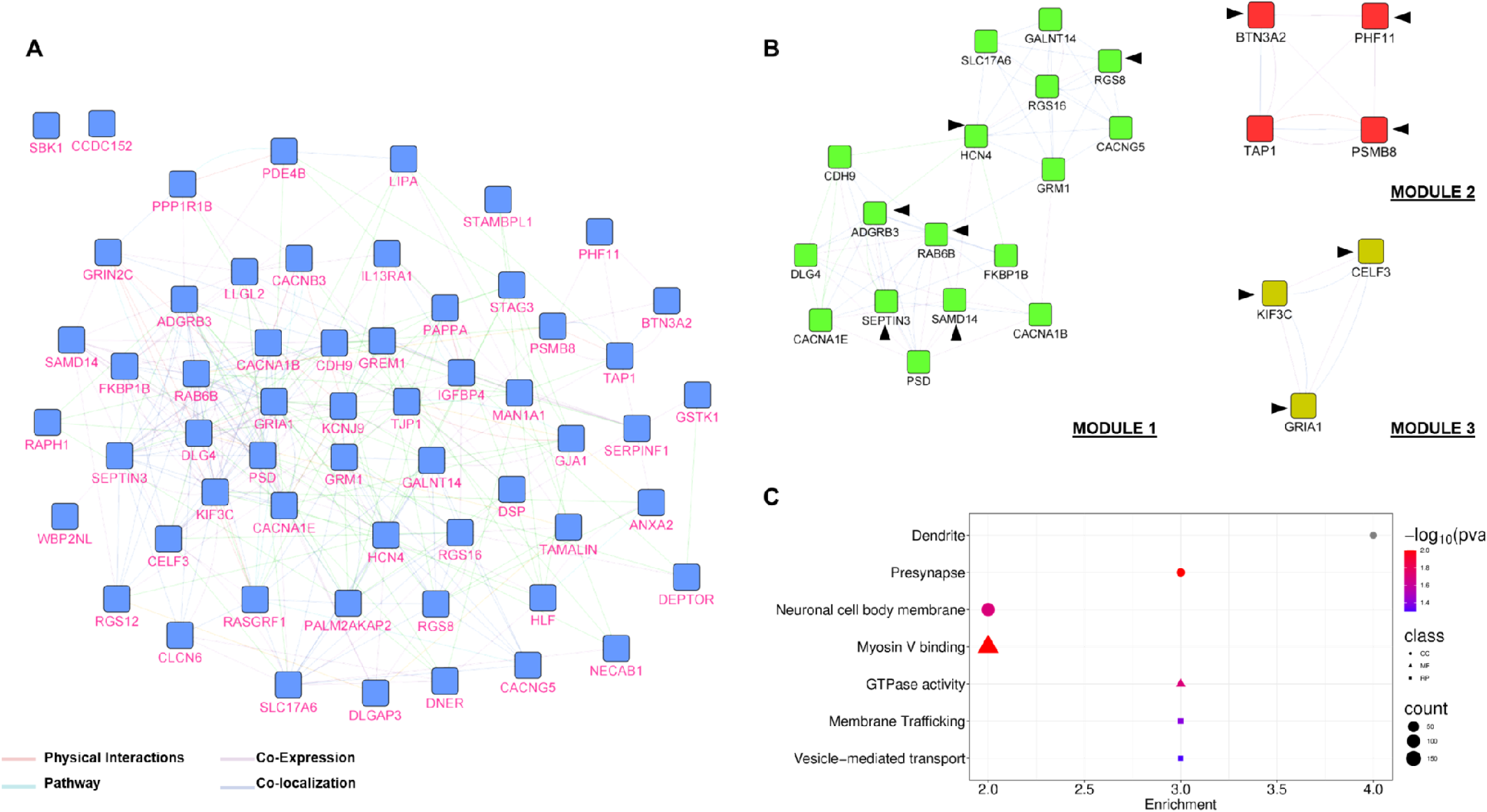
The Protein-Protein Interaction network and modular analysis of cDEGs. (A) The 40 cDEGs were filtered into the PPI network complex using the GENEMANIA plugin in the Cytoscape software. The network nodes include proteins encoded by the cDEGs and genes other than cDEGs. The edges represent their functional associations. (B) The three most significant modules within the PPI network were identified using the MCODE plugin in the Cytoscape software. The black arrowhead denotes cDEGs. (C) GO and Reactome pathway enrichment analysis of the hub genes.

### 3.3. Functional Enrichment

We performed GO analysis of the hub genes to identify key functional and pathway associations potentially driving DA neuron degeneration in familial and idiopathic PD. The analysis revealed two significant molecular functions (myosin V binding and GTPase activity) and three significant cellular components (dendrite, presynapse, and neuronal cell body membrane) (**Figure 2C**, **Table 3**). However, for biological processes, the GO analysis resulted in two non-significant biological processes (calcium-mediated signaling and neuron projection development). Pathway analysis through Reactome indicated that the hub genes involved membrane trafficking and vesicle-mediated transport (**Figure 2C**, **Table 3**). These findings suggest that hub genes, by regulating various critical functions, may play a pivotal role in the degeneration of DA neurons in familial and idiopathic PD.

**Table 3:** GO and Reactome Pathway Analysis of Hub Genes by DAVID.

| Category | Term | % | PValue | Genes | Fold Enrichment |
| --- | --- | --- | --- | --- | --- |
| GOTERM_BP_DIRECT | GO:0019722~calcium-mediated signaling | 16.67 | 0.06 | GRIA1, SAMD14 | 29.92 |
| GOTERM_BP_DIRECT | GO:0031175~neuron projection development | 16.67 | 0.06 | SAMD14, RAB6B | 27.58 |
| GOTERM_CC_DIRECT | GO:0030425~dendrite | 33.33 | 0.00 | HCN4, GRIA1, SAMD14, RGS8 | 14.81 |
| GOTERM_CC_DIRECT | GO:0098793~presynapse | 25 | 0.01 | GRIA1, SEPTIN3, RAB6B | 24.03 |
| GOTERM_CC_DIRECT | GO:0032809~neuronal cell body membrane | 16.67 | 0.02 | GRIA1, RGS8 | 114.81 |
| GOTERM_MF_DIRECT | GO:0031489~myosin V binding | 16.67 | 0.01 | GRIA1, RAB6B | 188.31 |
| GOTERM_MF_DIRECT | GO:0003924~GTPase activity | 25 | 0.02 | SEPTIN3, RGS8, RAB6B | 13.72 |
| REACTOME_PATHWAY | R-HSA-199991~Membrane Trafficking | 25 | 0.04 | GRIA1, KIF3C, RAB6B | 7.53 |
| REACTOME_PATHWAY | R-HSA-5653656~Vesicle-mediated transport | 25 | 0.05 | GRIA1, KIF3C, RAB6B | 7.09 |

### 3.4. Hub Gene-Disease Associations

The hub gene-disease network analysis identified four distinct subnetworks. Subnetwork 1 comprised the hub gene *PSMB8* and its association with 83 diseases (**Figure 3A**). Subnetwork 2 linked *GRIA1* to 13 diseases (**Figure 3B**), while Subnetwork 3 connected *HCN4* to 10 diseases (**Figure 3C**). The Subnetwork 4 showed an association between *PHF11* and two diseases (**Figure 3D**). Together, these subnetworks illustrate the varied disease associations of key hub genes, underscoring their potential roles in disease pathogenesis.

**Figure 3.**
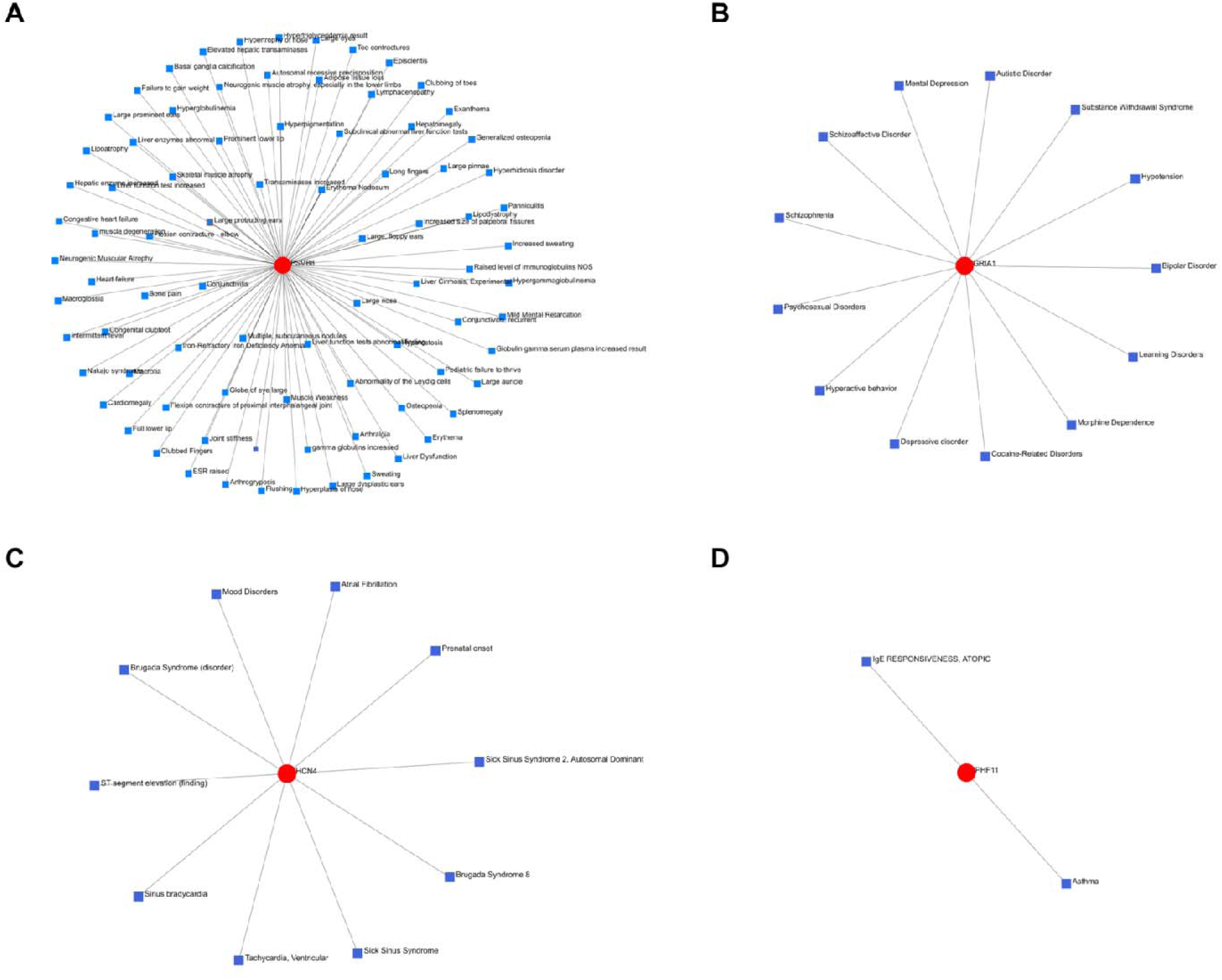
Hub Gene-Disease Interaction network created by Networkanalyst tool. Gene-Disease Interaction network of (A) PSMB8, (B) GRIA1, (C) HCN4, and (D) PHF11.

### 3.5. Validation of hub genes in the SH-SY5Y model of PD

We validated the expression patterns of the identified hub genes in an *in vitro* PD model using the neurotoxin MPTP-treated SH-SY5Y human neuroblastoma cells. qRT-PCR analysis demonstrated that, among the upregulated hub genes, *PSMB8* (2.03 ± 0.37; p < 0.04) and *BTN3A2* (1.66 ± 0.21; p < 0.04) exhibited significantly elevated expression levels in MPTP-treated SH-SY5Y cells compared to untreated controls (**Figure 4A**). Conversely, for the downregulated hub genes, qRT-PCR results revealed significantly decreased expression levels of *RGS8* (0.38 ± 0.14; p < 0.02), *CELF3* (0.39 ± 0.09; p < 0.003), *RAB6B* (0.74 ± 0.06; p < 0.02), *SEPTIN3* (0.57 ± 0.07; p < 0.005), and *HCN4* (0.42 ± 0.08; p < 0.002) in MPTP-treated cells relative to controls (**Figure 4B**). Overall, these results validate the differential expression of the identified hub genes in an *in vitro* PD model, reinforcing their potential significance in PD pathology.

**Figure 4.**
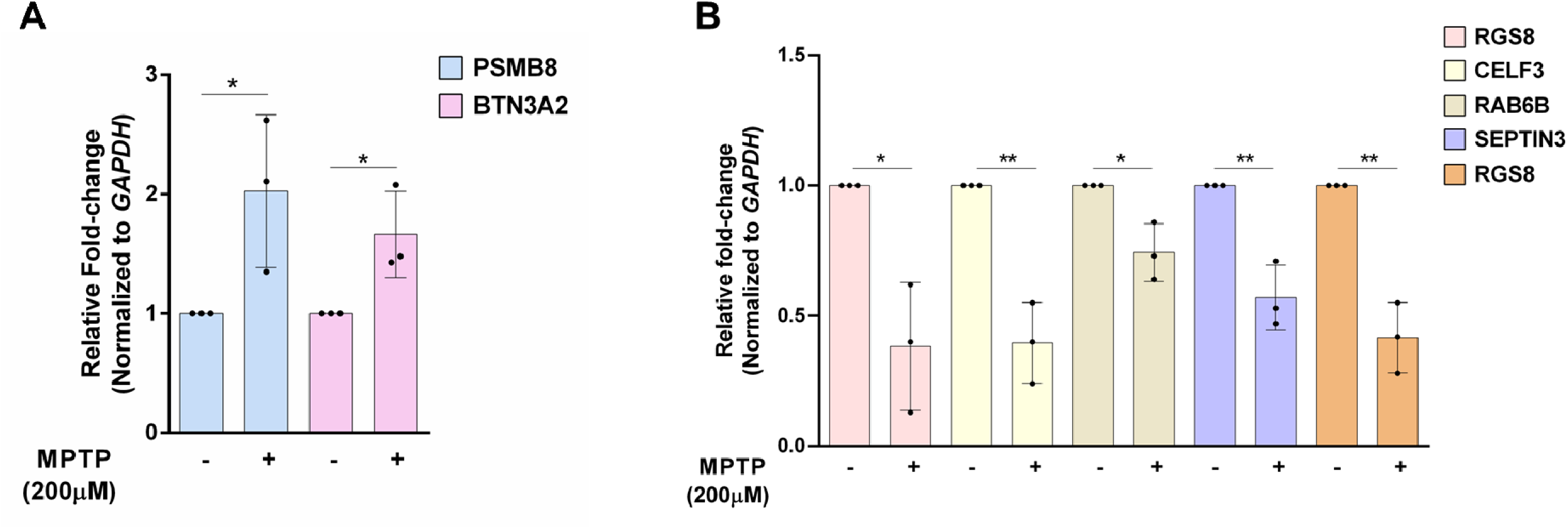
Expression profiles of hub genes in MPTP-treated SH-SY5Y cell model of PD versus untreated cells. Relative fold-change of (A) upregulated and (B) downregulated hub genes as validated by qPCR. mRNA for GAPDH was used for normalization. The data is represented as mean ± standard deviation. n = 3. * p<0.005, ** p<0.01; Unpaired t-test

### 3.6. *Rgs8* RNAi degenerates DA neurons in healthy *C. elegans*

The altered expression of the validated hub genes suggests they may play critical roles in maintaining DA neuron health. To test this hypothesis, we employed the *C. elegans* model, using the feeding RNAi method to investigate the functional significance of these hub genes. We first sought to determine which of the hub genes validated in MPTP-treated SH-SY5Y cells also exhibit altered expression in the nematode PD model UA44, which expresses human α-synuclein and GFP specifically in DA neurons. We used the BY250 strain as a healthy control, which expresses GFP in DA neurons but lacks α-synuclein expression. Both strains were fed control (empty vector) RNAi, and fluorescence microscopy performed at day 3 of adulthood revealed a significant reduction in fluorescence intensity in DA neurons of UA44 worms (9.47 ± 0.58, p < 0.0001) compared to BY250 worms (21.42 ± 1.14) (**Figure 5A and B**), confirming DA neuron degeneration in the PD model.

**Figure 5.**
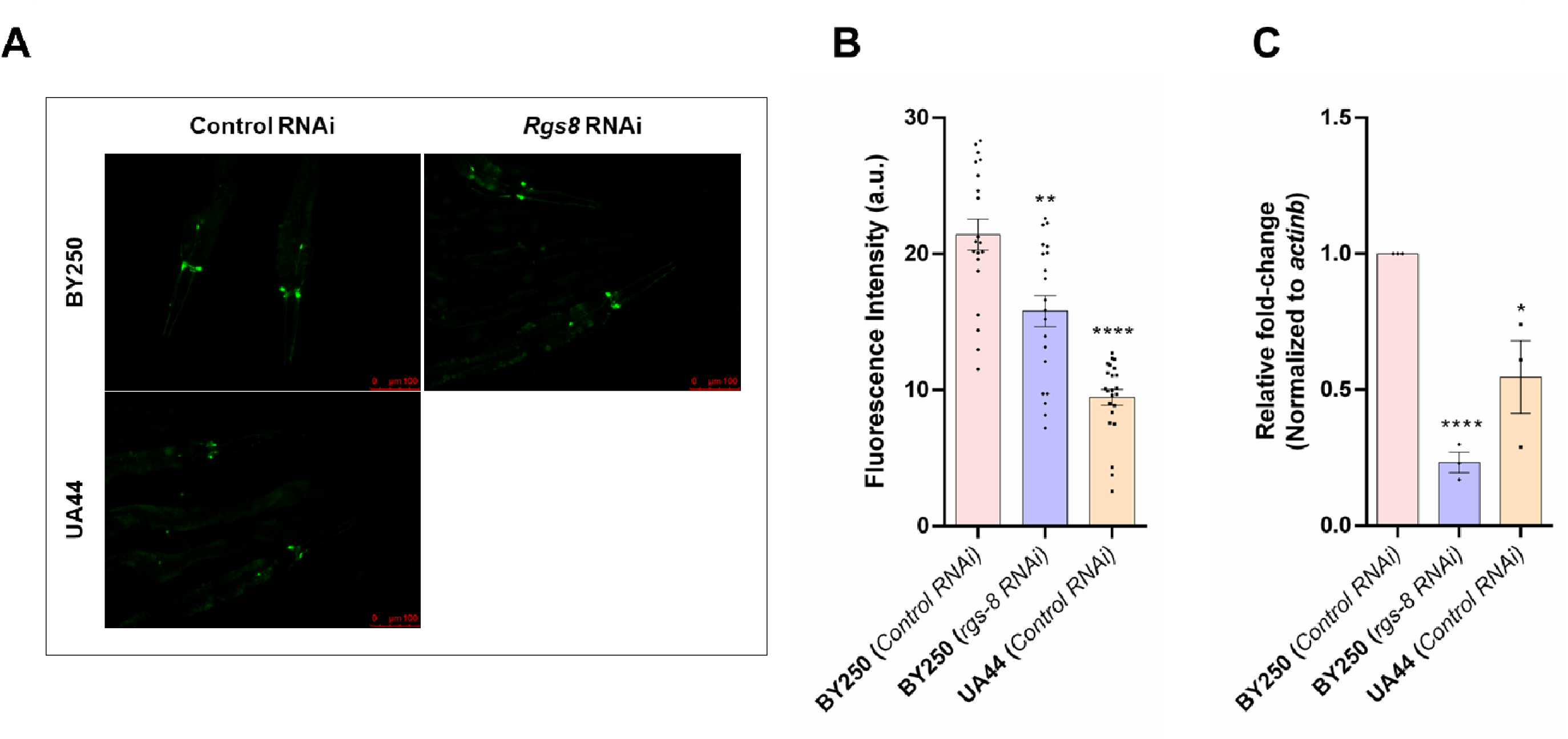
*Rgs8* RNAi affects DA neuronal health in *C. elegans*. (A) Fluorescence images of DA neurons within the head region of BY250 (Control RNAi), BY250 (*rgs8* RNAi), and UA44 (Control RNAi). (B) Graph showing fluorescence intensities of DA neurons in the head region as quantified by ImageJ software. The data is represented as mean ± standard deviation. n ≥ 20. ** p<0.01; **** p<0.0001; Unpaired t-test. (C) qRT-PCR data for *rgs8* mRNA in BY250 (Control RNAi), BY250 (*rgs8* RNAi) and UA44 (Control RNAi). mRNA for *actinb* was used for normalization. The data is represented as mean ± standard deviation. n=3. * p<0.05; **** p<0.0001; Unpaired t-test.

Subsequently, qRT-PCR analysis demonstrated a significant reduction in the mRNA levels of the *rgs8* homolog, *C29H12.3*, in UA44 worms (0.55 ± 0.13, p < 0.03) relative to BY250 controls (**Figure 5C**), suggesting a potential role for *C29H12.3/rgs8* in DA neurodegeneration. To further investigate this, we performed RNAi-mediated knockdown of *C29H12.3/rgs8* in BY250 worms and evaluated its impact on DA neuron integrity. qRT-PCR confirmed efficient knockdown, with a significant reduction in *C29H12.3/rgs8* mRNA levels (0.23 ± 0.04, p < 0.0001) in *C29H12.3/rgs8* RNAi-fed worms compared to control RNAi-fed worms (**Figure 5C**). Consistent with a role in neuronal maintenance, fluorescence microscopy at day 3 of adulthood revealed a significant decrease in fluorescence intensity in DA neurons of *C29H12.3/rgs8* RNAi-treated BY250 worms (15.81 ± 1.15, p < 0.002) compared to controls (21.42 ± 1.14) (**Figure 5A and B**). These results suggest that *C29H12.3/rgs8* is essential for maintaining DA neuron integrity and may contribute to the molecular mechanisms underlying neurodegeneration in PD.

## 4. Discussion

This study offers valuable insights into the shared molecular mechanisms underlying DA neuron degeneration in familial and idiopathic PD forms. Previous research has demonstrated the utility of reanalyzing publicly available transcriptomic datasets to uncover novel biological insights ^71–78^. In this study, we integrated RNA sequencing data from three independent datasets, each representing distinct genetic and pathological contexts of PD. Through this approach, we identified 40 cDEGs, including 12 hub genes, which may play key roles in the pathogenesis of PD. Functional enrichment analysis revealed that hub genes participate in pathways essential for maintaining neuronal health and synaptic integrity—disrupted processes in PD.

Although the datasets represent samples with differing genetic backgrounds—potentially influencing disease onset and progression—our aim was to identify converging molecular alterations that may represent shared pathogenic mechanisms across PD subtypes. The identification and subsequent experimental validation of these genes underscore the existence of common molecular disruptions in familial and idiopathic PD, highlighting potential targets for therapeutic intervention.

Notably, several of the cDEGs identified in our analysis, including *PPP1R1B*, *DEPTOR*, *DLGAP3*, *GJA1*, *IL13RA1*, *LIPA*, *NECAB1*, and *PSMB8*, have previously been reported to exhibit altered expression in PD models and human PD brain tissues ^79–104^. Although these prior studies primarily focused on idiopathic PD, our findings extend the relevance of these genes by demonstrating their dysregulation across both idiopathic and familial PD datasets, suggesting that their involvement in DA neuron degeneration may represent a common molecular signature of PD independent of genetic origin.

The hub genes, *PSMB8, GRIA1,* and *RGS8,* exhibit expression changes consistent with previous reports, supporting their potential relevance to PD pathophysiology ^105–107^. In addition, our study demonstrates that *rgs8* mRNA levels are significantly reduced not only in the MPTP-treated SH-SY5Y cell model of PD but also in the transgenic *C. elegans* PD model, complementing earlier findings of reduced *rgs8* expression in the rat α-synuclein preformed fibril model ^107^. Notably, this is the first study to show, using the *C. elegans* model, that *rgs8* downregulation leads to DA neurodegeneration. RGS8 (Regulator of G Protein Signaling 8) is known to regulate G protein-coupled receptor (GPCR) signaling by acting as a GTPase-activating protein ^108^. Given that GPCRs are essential to dopamine signaling, RGS8 may be crucial in maintaining DA neuron health ^109^. Future studies should aim to elucidate the molecular mechanisms by which *rgs8* regulates DA neuron integrity and its potential contribution to PD progression.

The other hub genes are linked to DA neuron dysfunction in PD for the first time in this study (**Table 4**). For example, *BTN3A2*, a member of the butyrophilin subfamily, is primarily known for its role in modulating immune responses ^110^. Since neuroinflammation is a well-established contributor to PD pathogenesis, the upregulation of BTN3A2 observed in PD-DA neurons suggests a potential link between peripheral immune signaling and central neurodegenerative processes in idiopathic and familial PD ^111^. Similarly, *HCN4* (Hyperpolarization-Activated Cyclic Nucleotide-Gated Potassium Channel 4), which modulates neuronal excitability via potassium channel regulation, also emerged as a downregulated hub gene. Alterations in *HCN4* expression may impair neuronal membrane potentials and contribute to excitotoxicity—a well-documented mechanism of neurodegeneration in PD ^112,113^. Together, these findings highlight the previously unrecognized involvement of several genes in DA neuron degeneration and suggest that these hub genes may represent critical molecular nodes in PD pathogenesis. Further mechanistic studies are needed to elucidate their functional roles and therapeutic potential.

**Table 4:** Potential Role of Hub Genes in DA neurodegeneration.

| <u>S.no.</u> | <u>Hub Gene</u> | <u>Protein Encoded</u> | <u>Expression in GSE90469, GSE169755, and GSE181029 (PD vs. control)</u> | <u>Reported Cellular Function</u> | <u>Reported PD-association</u> | <u>Potential Role in DA-neurodegeneration</u> |
| --- | --- | --- | --- | --- | --- | --- |
| 1 | *PSMB8* | Proteasome subunit beta type-8 | Upregulated | Part of 20S proteasome complex <sup>105</sup> | Yes <sup>105</sup> | May affect the clearance of misfolded or aggregated proteins |
| 2 | *GRIA1* | Glutamate receptor 1 | Downregulated | Subunit of the AMPA-type glutamate receptor <sup>118</sup> | Yes <sup>106</sup> | May disrupt synaptic signaling and exacerbate excitotoxic stress |
| 3 | *RGS8* | Regulator of G-protein signaling 8 | Downregulated | Regulate G protein-coupled receptor <sup>119</sup> | Yes <sup>107</sup> | May exacerbate immune-mediated neurotoxicity |
| 4 | *PHF11* | Plant Homeodomain Finger 11 | Upregulated | Regulate transcription and inflammatory signaling pathways <sup>120</sup> | No | May promote a pro-inflammatory environment and alter stress response pathways |
| 5 | *RAB6B* | Ras-related protein Rab-6B | Downregulated | Regulate vesicle trafficking and transport within the Golgi complex <sup>121</sup> | No | May impair the clearance of aggregated proteins |
| 6 | *CELF3* | CUGBP Elav-Like Family Member 3 | Downregulated | Regulate RNA processing, including cotranscriptional and post-transcriptional events <sup>122</sup> | No | May impair gene regulation |
| 7 | *ADGRB3* | Adhesion G Protein-Coupled Receptor B3 | Downregulated | Regulate synapse formation and maintenance <sup>123</sup> | No | May impair synaptic function |
| 8 | *BTN3A2* | Butyrophilin Subfamily 3 Member A2 | Upregulated | Regulate adaptive immune response and immune checkpoint regulation <sup>124</sup> | No | May mediate immune-driven neuronal injury |
| 9 | *HCN4* | Hyperpolarization Activated Cyclic Nucleotide Gated Potassium Channel 4 | Downregulated | Regulate neuronal excitability <sup>125</sup> | No | May disrupt DA neuronal pacemaking and calcium homeostasis. |
| 10 | *KIF3C* | Kinesin family member 3C | Downregulated | Regulate neuronal development and axon growth and regeneration<sup>126</sup> | No | May exacerbate axonal pathology |
| 11 | *SEPTIN3* | Septin 3 | Downregulated | Regulate memory and long-term potentiation<sup>127</sup> | No | May disrupt synaptic homeostasis |
| 12 | *SAMD14* | Sterile Alpha Motif Domain Containing 14 | Downregulated | Regulate stress response pathways<sup>128</sup> | No | May compromise neuronal stress responses |

Genes play a central role in regulating a wide array of cellular functions, and disruptions in their expression can perturb essential biological pathways, thereby contributing to the development and progression of various diseases. Our gene-disease network analysis revealed one of the hub genes, *GRIA1,* with established associations with bipolar disorder—a neuropsychiatric condition marked by altered DA signaling during manic and depressive phases ^114,115^. Notably, emerging evidence suggests that individuals with bipolar disorder may be at an elevated risk of developing PD later in life ^116^. Although polymorphisms in *GRIA1* have been linked to psychotic subtypes of bipolar disorder, their specific role in PD pathogenesis remains undefined and merits further investigation ^117^. Uncovering shared genetic contributors such as *GRIA1* across distinct neurological disorders holds promise for developing therapeutic strategies that target convergent molecular mechanisms.

Collectively, these findings strengthen the case for shared molecular pathways across familial and idiopathic PD forms and provide a foundation for future research into therapeutic targeting of hub genes. Given the limitations of existing PD treatments—largely symptomatic and failing to halt disease progression—targeting common pathogenic mechanisms may offer broader clinical benefits.

However, several limitations of this study warrant consideration. First, the reliance on transcriptomic data captures only one layer of gene regulation, and additional studies incorporating proteomic, epigenetic, and metabolomic data could provide a more comprehensive view. Second, while SH-SY5Y and *C. elegans* models are powerful for validation, they do not fully replicate the complexity of human DA neurons and brain architecture. Further *in vivo* validation in mammalian models, such as mice or non-human primates, is necessary to confirm the translational relevance of the findings. Finally, the functional roles of many hub genes remain poorly understood, and in-depth mechanistic studies will be essential to elucidate their contributions to DA neuron vulnerability.

Nevertheless, our study provides valuable insights into the DA neuron-specific shared molecular mechanisms underlying familial and idiopathic PD. These findings expand the landscape of molecular players in PD pathogenesis, providing new hypotheses for experimental validation. We hope that the findings from our study will inspire future experimental validation, fostering new insights into shared molecular mechanisms and uncovering novel therapeutic targets to deepen our understanding of PD.

## Abbreviations

PD: Parkinson’s Disease
DA: Dopaminergic Neurons
SH-SY5Y: Human Neuroblastoma Cell Line
SN: Substantia Nigra
MPTP: 1-Methyl-4-Phenyl-1,2,3,6-Tetrahydropyridine
qRT-PCR: Quantitative Reverse Transcription Polymerase Chain Reaction
PPI: Protein-Protein Interaction
GO: Gene Ontology
GEO: Gene Expression Omnibus
DEGs: Differentially Expressed Genes
cDEGs: Common Differentially Expressed Genes
iPSCs: Induced Pluripotent Stem Cells
GWAS: Genome-Wide Association Study
BTN3A2: Butyrophilin Subfamily 3 Member A2
GRIA1: Glutamate Ionotropic Receptor AMPA Type Subunit 1
PSMB8: Proteasome Subunit Beta Type-8
HCN4: Hyperpolarization-Activated Cyclic Nucleotide-Gated Potassium Channel 4
RGS8: Regulator of G Protein Signaling 8
CELF3: CUGBP Elav-Like Family Member 3
SEPTIN3: Septin 3
GPCR: G Protein-Coupled Receptor
PHF11: Plant Homeodomain Finger Protein 11
PARK2: Parkinsonism Associated Deglycase (Parkin)
LRRK2: Leucine-Rich Repeat Kinase 2
PINK1: PTEN-Induced Kinase 1
DJ-1: Protein Deglycase DJ-1
SNCA: Synuclein Alpha
GBA: Glucocerebrosidase
DisGeNET: Disease Gene Association Network
DAVID: Database for Annotation, Visualization, and Integrated Discovery
MCODE: Molecular Complex Detection
DMSO: Dimethyl Sulfoxide
IPTG: Isopropyl β-D-1-Thiogalactopyranoside
LB: Luria-Bertani Medium
NGM: Nematode Growth Medium
RNAi: RNA Interference
FC: Fold Change
L1: First Larval Stage

## Acknowledgments

S is supported by Senior Research Fellowship from CSIR. GC is supported by Junior Research Fellowship UGC. We thank the Director of CSIR-CDRI for providing the necessary research facilities. We sincerely thank the faculty members of the Division of Neuroscience and Ageing Biology, CSIR-CDRI, for providing access to the cell culture facility and the RT-PCR machine (Biorad-CFX96). We would also like to acknowledge Dr. Aamir Nazir, Division of Toxicology & Experimental Medicine, CSIR-CDRI, for sharing *C29H12.3/rgs8* RNAi and providing access to the fluorescence microscope (Carl Zeiss).

## CRediT authorship contribution statement

**S-** Investigation; Data curation; Methodology; Writing—original draft; Writing—review and editing.

**GC**- Investigation; Methodology; Writing—original draft.

**BM-** Resources; Investigation. Writing - review & editing.

**SV**- Conceptualization; Investigation; Data curation; Formal analysis; Methodology; Project administration; Resources; Supervision; Validation; Visualization; Writing - original draft; and Writing - review & editing.

## Data Availability

The datasets used in this study are available in the GEO database [https://www.ncbi.nlm.nih. gov/geo/]. Details about the corresponding accession numbers are provided within the article.

---

**Supplementary Table 1:**
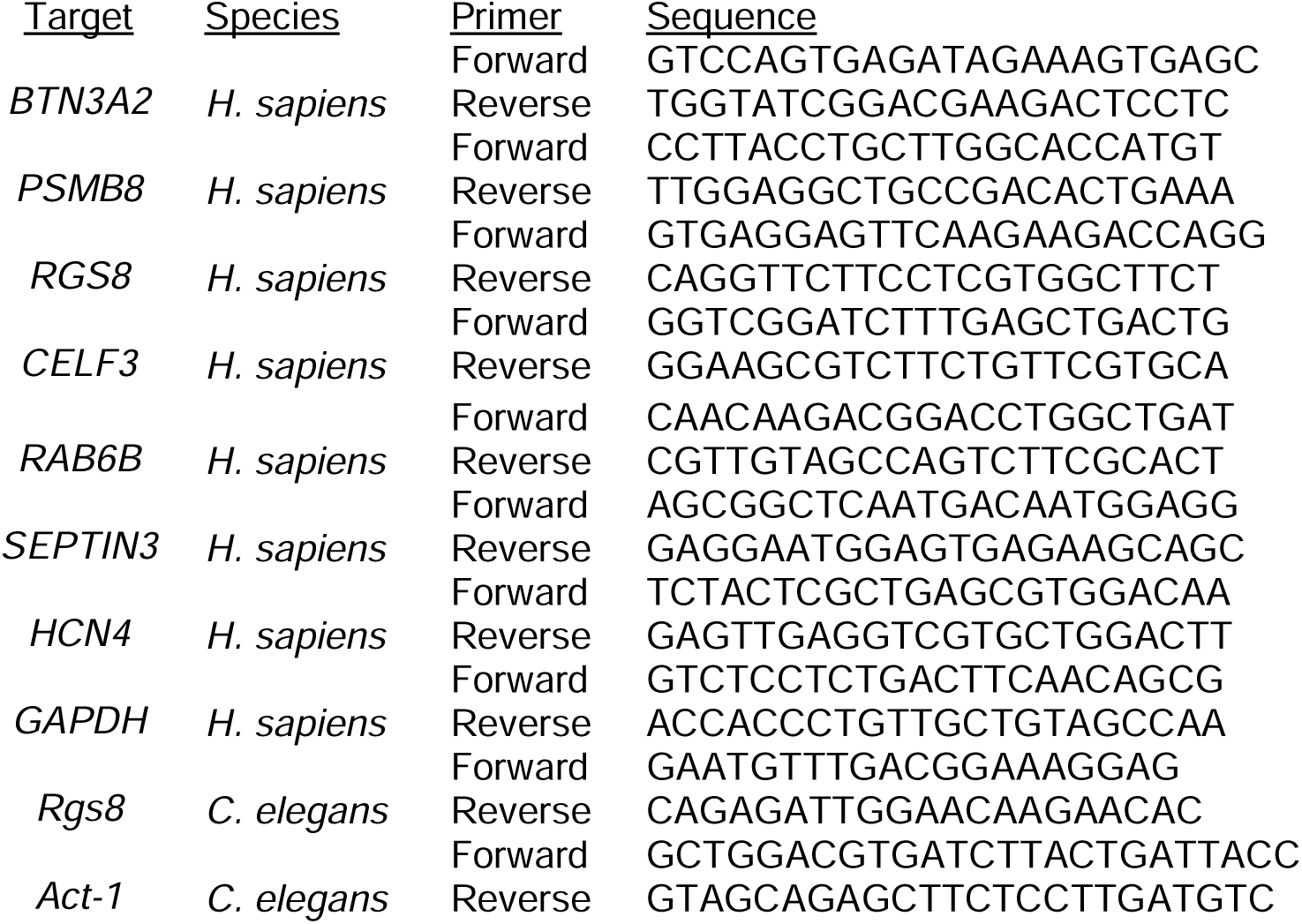
Sequence of Primers used for qRT-PCR.

